# Cryopreservation of human mucosal tissues

**DOI:** 10.1101/262519

**Authors:** Sean M. Hughes, April L. Ferre, Sarah E. Yandura, Cory Shetler, Chris A. R. Baker, Fernanda Calienes, Claire N. Levy, Rena D. Astronomo, Zhiquan Shu, Gretchen M. Lentz, Michael Fialkow, Anna C. Kirby, M. Juliana McElrath, Elizabeth Sinclair, Lisa C. Rohan, Peter L. Anderson, Barbara L. Shacklett, Charlene S. Dezzutti, Dayong Gao, Florian Hladik

## Abstract

**BACKGROUND:** Cryopreservation of leukocytes isolated from the cervicovaginal and colorectal mucosa is useful for the study of cellular immunity (see Hughes SM et al. PLOS ONE 2016). However, some questions about mucosal biology and sexually transmitted infections are better addressed with intact mucosal tissue, for which there is no standard cryopreservation protocol.

**METHODS AND FINDINGS:** To find an optimal preservation protocol for mucosal tissues, we tested slow cooling (1%C/min) with 10% dimethylsulfoxide (designated “cryopreservation”) and fast cooling (plunge in liquid nitrogen) with 20% dimethylsulfoxide and 20% ethylene glycol (“vitrification”). We compared fresh and preserved human cervicovaginal and colorectal tissues in a range of assays, including metabolic activity, human immunodeficiency virus infection, cell phenotype, tissue structure by hematoxylin-and-eosin staining, cell number and viability, production of cytokines, and microbicide drug concentrations. Metabolic activity, HIV infectability, and tissue structure were similar in cryopreserved and vitrified vaginal tissues. However, vitrification led to poor cell recovery from the colorectal mucosa, with 90% fewer cells recovered after isolation from vitrified colorectal tissues than from cryopreserved. HIV infection rates were similar for fresh and cryopreserved ectocervical tissues, whereas cryopreserved colorectal tissues were less easily infected than fresh tissues (hazard ratio 0.7 [95% confidence interval 0.4, 1.2]). Finally, we compared isolation of cells before and after cryopreservation. Cell recoveries were higher when cells were isolated after freezing and thawing (71% [59-84%]) than before (50% [38-62%]). Cellular function was similar to fresh tissue in both cases. Microbicide drug concentrations were lower in cryopreserved explants compared to fresh ones.

**CONCLUSIONS:** Cryopreservation of intact cervicovaginal and colorectal tissues with dimethylsulfoxide works well in a range of assays, while the utility of vitrification is more limited. Cell yields are higher from cryopreserved intact tissue pieces than from thawed cryopreserved single cell suspensions isolated before freezing, but T cell functions are similar.

## Introduction

The availability of mucosal tissue specimens is crucial for the study of mucosal biology and immunity. Currently, working with mucosal specimens is complicated by the difficulty of storing tissues long-term while maintaining good viability. We have recently reported methods for the cryopreservation of leukocytes isolated from mucosal tissues [1], but limited and conflicting data are available for intact tissues [2–5].

A protocol for preserving tissue specimens with good viability and function would enable their collection and analysis in clinical trials involving STIs such as human immunodeficiency virus (HIV) or herpes simplex virus. These trials are conducted at sites around the world and samples must be transported to analysis laboratories. Because there is no standard preservation protocol, assays requiring viable tissue cannot currently be done. Instead, samples are typically preserved according to their intended use. For example, samples intended for measurement of drug concentration might be snap frozen, while samples intended for immunohistochemistry might be preserved in formalin. The ability to preserve tissues with good viability would not only allow functional assays to be performed, but also might allow samples to be stored without pre-determining the type of assay to be performed on them. Storing viable samples could thereby give the investigator more options, as the viable samples could be thawed and then preserved according to the needs of the assay later on.

We tested two methods for preservation: cryopreservation and vitrification. Cryopreservation involves freezing cells or tissues in the presence of relatively low concentrations of a cryoprotective agent (such as 10% dimethylsulfoxide [DMSO]), typically at a slow rate (such as, −1°C/min). Cryopreservation is widely used for many cell types. Vitrification involves higher concentrations of cryoprotective agents (around 40%) and freezing as fast as possible (such as directly submerging in liquid nitrogen). Successful vitrification of intact tissues similar to the colorectal and cervicovaginal mucosa has been reported for skin [6–10], testicles [11,12], umbilical cord [13], trachea [14], and oral mucosa [15]. Vitrification has been widely adopted for in vitro fertilization because of similar or even improved clinical outcomes as compared to fresh and cryopreserved embryos or oocytes [16]. However, there are few reports of successful cryopreservation or vitrification of intact colorectal or cervicovaginal mucosa.

We therefore developed cryopreservation and vitrification protocols for colorectal and cervicovaginal mucosa by sequentially testing procedural variations, informed by studies of the fundamental cryobiological properties of these tissues [17,18]. Once we developed two final protocols, one for vitrification and one for cryopreservation, we tested these protocols more thoroughly. While we show that both procedures work well in several assays with vaginal tissues, we found that cryopreservation was dramatically superior for isolation of single cells from colorectal tissues and, moreover, worked well for a range of assays with vaginal, ectocervical, and colorectal tissues. As cryopreservation was superior in some assays, equivalent in others, and simpler to perform than vitrification, we recommend it and provide a detailed protocol for tissue cryopreservation in supporting **S1 Text**. Finally, we compared two cryopreservation procedures for isolated cells: cryopreservation of intact tissue pieces with cell isolation after thawing or cryopreservation of isolated cells. Cytokine production was similar to that of fresh samples with both procedures, but yield and viability were higher for cryopreservation of intact tissue pieces, so we recommend that method.

## Methods

### Sample collection

Two types of tissue were used for this study: explants, which were derived from discarded surgical tissues dissected in the laboratory, and biopsies, which were obtained directly from the volunteer through flexible sigmoidoscopy. Vaginal tissues were collected under a waiver of consent from the University of Washington Medical Center (IRB #1167) during vaginal repair surgeries. Colorectal biopsies were obtained by flexible sigmoidoscopy at San Francisco General Hospital (UCSF IRB #10-01218 and 10-00263). Normal human ectocervical (IRB #0503103) and colorectal (IRB #0602024) tissues were acquired from pre-menopausal women undergoing hysterectomy or persons undergoing colorectal surgery for non-inflammatory conditions through the University of Pittsburgh, Health Sciences Tissue Bank. Normal human ectocervical tissue was also purchased from National Disease Research Interchange (NDRI) (http://ndriresource.org/) and shipped overnight on ice. All participants provided written informed consent. Vaginal tissues were placed into cold phosphate-buffered saline and transported on ice to the laboratory. Colorectal biopsies were collected in Roswell Park Memorial Institute-1640 medium (RPMI) containing 15% fetal bovine serum with 100 units/mL penicillin, 100 μg/mL streptomycin, and 2 mM L-glutamine. Ectocervical and surgical colorectal tissues were placed in L-15 medium supplemented with 10% fetal bovine serum, 100 µg/ml streptomycin, 100 U/ml penicillin, and 2.5 µg/ml Amphotericin B for transport. All tissues were transported to the laboratory on ice and processed within 4 hours of surgery or flexible sigmoidoscopy, except for those ectocervical tissues which were purchased from the NDRI and were shipped overnight on ice.

### Assay sites

This was a multi-site study, with different assays performed on different types of tissue at each site. All studies with vaginal explants (alamar blue, hematoxylin-and-eosin staining, and HIV infection) were performed in Seattle, WA with tissues obtained in the same location. All studies with ectocervical explants (HIV infection, drug levels) as well as the HIV infections of colorectal explants were performed in Pittsburgh, PA with tissues obtained in the same location or from the NDRI. Phenotyping of cells from digested colorectal biopsies was performed in San Francisco, CA with tissues obtained in the same location. Stimulation and intracellular cytokine staining of cells from digested colorectal biopsies was performed in Davis, CA with tissues shipped from San Francisco.

### Surgical tissue preparation

Vaginal tissues were placed in a petri dish for dissection and covered with cold phosphate-buffered saline (PBS). They were trimmed to a thickness of 2-4 mm. Uniform cylindrical explants 2 mm in diameter were then obtained with punch biopsies (Miltex, York, PA, USA). Explants were transferred to fresh, cold phosphate-buffered saline and used for experiments. For ectocervical and colorectal tissue, 5 mm explants were made using a dermal punch. As tissue size can influence assay results, multiple explants or biopsies were used for each experiment and we distributed tissue pieces to have a consistent range of sizes across conditions.

### Chemicals and cryoprotective agents

Dimethylsulfoxide (DMSO) and ethylene glycol (EG) were obtained from Sigma-Aldrich (St. Louis, MO, USA). Trehalose and copper foil were obtained from Merck Millipore (Billerica, MA, USA). 10% neutral buffered formalin was obtained from Electron Microscopy Sciences (Hatfield, PA, USA).

### Cryopreservation and vitrification procedures

For cryopreservation, tissues were transferred into cryovials containing 200 µL 10% DMSO in fetal bovine serum. Cryovials were sealed and frozen to −80°C in Mr. Frosty controlled cooling rate devices (Thermo Fisher Scientific, Waltham, MA, USA). Cryovials were then transferred to the vapor phase of a liquid nitrogen freezer within one week. To thaw, cryovials were placed in a 37°C water bath and agitated until only a pea-sized piece of ice remained. Thawed tissues were transferred with forceps or sterile transfer pipets to 10-20 mL of room temperature cell culture medium and allowed to equilibrate for 10 min before further use. Single explants were cryopreserved for ectocervical, vaginal, and colorectal surgical tissues, while up to 15 colorectal biopsies were cryopreserved per cryovial with 1 mL cryopreservation medium. A detailed cryopreservation protocol is available in **S1 Text** and at https://dx.doi.org/10.17504/protocols.io.p5adq2e.

For vitrification, tissues were placed in half-concentration vitrification medium (10% DMSO with 10% EG in a solution of phosphate-buffered saline containing 20% fetal bovine serum) for five minutes and then a full-concentration (20% DMSO and 20% EG) solution for five minutes, both at room temperature. They were blotted dry and placed on a vitrification device (a piece of aluminum foil or plastic straw [Cryo Bio System, Maple Grove, MN, USA]). Single explants were always used for vaginal tissue, while up to 5 colorectal biopsies were vitrified per device. The devices were plunged directly into liquid nitrogen and held there until bubbling subsided. They were then transferred into pre-cooled cryovials and stored in the vapor phase of a liquid nitrogen freezer. To thaw, cryovials were opened while still on liquid nitrogen, and the foil was rapidly transferred into room temperature thawing medium (10% DMSO and 10% EG in cell culture medium) and agitated until the tissue piece thawed enough to fall off into the media. After 10 min of equilibration, tissues were moved to cell culture medium for an additional 10 min. A detailed vitrification protocol is available in **S1 Text** and at https://dx.doi.org/10.17504/protocols.io.p6adrae.

### Measurement of metabolic activity with alamar blue

Explants were frozen and thawed the same day to be synchronized with fresh explant controls. Specifically, cryopreserved explants were frozen at −80C in a controlled rate cooling device for 90 min (sufficient to freeze to −80C) and then placed in liquid nitrogen for a few minutes. Vitrified explants were frozen and held on liquid nitrogen until the cryopreserved explants were ready. Fresh explants were stored in culture during this time. After thawing the cryopreserved and vitrified tissues, all explants were cultured overnight in cell culture medium (RPMI containing 10% fetal bovine serum with 100 units/mL penicillin and 100 µg/mL streptomycin) in a humidified, 5% CO_2_ incubator at 37°C. The next morning, alamar blue (resazurin, Invitrogen, Carlsbad, CA, USA) was added to achieve a ratio of 1:10 alamar blue to culture media and culture was continued for 5 hours. To measure alamar blue fluorescence, media were mixed by pipetting and 90 µL aliquots were transferred to white-bottomed plates and measured on a SpectraMax M2 plate reader (Molecular Devices, Sunnyvale, CA, USA) with 544 nm excitation and 590 nm emission.

### HIV infections

After thawing, one explant was cultured per well of a 96-well plate and challenged with HIV as described previously [19]. Briefly, explants were activated overnight with 0.6 µg/mL phytohemagglutinin (PHA), 500 IU/mL interleukin-2 (IL-2), 25 ng/mL interleukin-7 (IL-7), and 5 ng/mL interleukin-15 (IL-15) (Peprotech, Rocky Hill, NJ, USA). The next day, the activation media was replaced with media containing the cytokines without PHA and explants were challenged with 200,000 IU per well of nanoluciferase-secreting JR-CSF HIV-1. The next day, explants were washed five times to remove the viral inoculum. Explants were then maintained in media containing IL-2, IL-7, and IL-15 for three weeks, with media change and supernatant collection every 2-3 days. Supernatants were stored at −80°C.

After completion of culture, nanoluciferase levels were measured in the supernatants with the Nano-Glo luciferase assay system (Promega, Madison, WI, USA). Supernatants were equilibrated to room temperature for at least 30 min. The substrate was prepared according to the manufacturer’s instructions and 20 µL of supernatant was mixed with 20 µL of substrate. After five minutes, luminescence was measured on the SpectraMax M2 plate reader.

Individual colorectal explants were placed on medium-soaked gelfoam rafts, while ectocervical explants were placed directly into a well of a 48-well plate with 1 mL of culture medium. Cultures were maintained at 37°C in a 5% CO_2_ atmosphere[20–24]. Each treatment was performed with duplicate explants. Fresh control explants were prepared on day of surgery in duplicate and treated with HIV-1_BaL_ (5×10^4^ TCID_50_ for ectocervix and 1×10^4^ TCID_50_ for colorectal). Cryopreserved explants were thawed and then processed in a similar manner as the fresh explants. The explants remained in culture for 11 to 14 days with fresh basolateral medium replenished every 3 to 4 days. The culture supernatant was stored at −80°C. The stored supernatant was tested for HIV-1 replication using the p24gag ELISA (Perkin-Elmer Life and Analytical Sciences, Inc., Waltham, MA).

### Hematoxylin and eosin

Vaginal explants were prepared as described above from five donors. One explant was placed directly into 10% neutral buffered formalin for one week and then transferred to 70% ethanol. The second explant was cryopreserved and the third explant was vitrified. Frozen samples were stored in a liquid nitrogen freezer for 1-2 months. After thawing, the samples were transferred to 10% neutral buffered formalin and left for one week. After one week, the explants were transferred to 70% ethanol.

Explants were processed and embedded into paraffin blocks and hematoxylin and eosin stains were performed on two slices from each block. The slides were imaged at 40X on an Aperio ScanScope (Leica Biosystems, Nußloch, Rhein-Neckar-Kreis, Germany).

### Colorectal tissue digestions and cell purification

For comparison of vitrification and cryopreservation, colorectal biopsies were digested after thawing with collagenase to yield single cell suspensions [25]. Biopsies were transferred to a 50 mL conical tube containing 12 mL of RPMI with 5% fetal bovine serum, 10 μg/mL DNase I (Sigma) and 700 units/mL collagenase type II (Sigma), and agitated at 37°C for 30 min. After incubation, a blunt-ended 16-gauge needle was used to mechanically break down the tissue and then the suspension was filtered through a 70 μm cell strainer to separate free cells from undigested biopsies. Undigested tissue was subjected to another 30 min round of digestion with 10 mL of fresh collagenase/DNase solution. Digested fractions were combined and washed in fresh RPMI with 5% fetal bovine serum and 10 μg/mL DNase I. The cell suspension was resuspended in a solution of 44% Percoll (GE Healthcare, Little Chalfont, United Kingdom) in RPMI and overlaid onto a solution of 66% Percoll in phosphate-buffered saline. The cells were centrifuged at 800g for 15 min to isolate mucosal mononuclear cells from epithelial cells and residual mucus. The mononuclear cell layer at the interface was removed with a sterile serological pipet and washed twice in PBS with 2% FBS and 10 mM ethylenediaminetetraacetic acid (EDTA).

For comparison of fresh, cryopreserved cell suspensions, and cryopreserved intact tissue, colorectal biopsies were digested in a similar manner, except that a different digestion medium was used (50 μg/mL Liberase DL [catalog #05 466 202 001, Roche Diagnostics, Mannheim, Germany] in RPMI with 100 units/mL penicillin, 100 µg/mL streptomycin, 2 mM L-glutamine, and 25 mM HEPES buffer), the digestions were repeated up to four times, and no subsequent Percoll gradient step was included. Cells frozen in suspension for these experiments were frozen as described previously [1]. Isolated cells to be used for intracellular staining were rested overnight in cell culture media containing 0.5 mg/mL piperacillin/tazobactam (Zosyn, Wyeth-Ayerst, Princeton, NJ, USA.)

### Colorectal cell counting and phenotyping

After Percoll isolation, a small aliquot of cells was resuspended in ViaCount (Millipore) and analyzed on an Accuri C6 (BD Biosciences, San Jose, CA, USA) benchtop, volumetric cytometer to assess cell count and viability.

One million cells were plated per well in 96-well V-bottom plates, resuspended in PBS, and stained for 15 min with live/dead fixable aqua dead cell stain (Thermo Fisher Scientific). The cells were then incubated with purified human IgG to block Fc receptors for 10 min at 4°C. After washing with PBS containing 2% fetal bovine serum and 2 mM EDTA, cells were stained a panel of phenotyping antibodies, including CD45, CD3, CD4, CD8, CD66b, CD33, and CD13. Complete antibody identification, including suppliers, catalog numbers, clones, and fluorochromes, is given in **S1 Table**. Cells were stained for 15 minutes at 4°C, washed, fixed in 1% paraformaldehyde in PBS, and acquired on an LSRII flow cytometer (BD Biosciences).

### Colorectal *ex vivo* stimulation and intracellular staining

Intracellular cytokine staining was performed as previously described [1,25]. Briefly, cells were stimulated with one of the following: a pool of peptides from cytomegalovirus, Epstein-Barr virus, and influenza (AnaSpec, Fremont, CA, USA); an HIV-1 Gag p55 peptide pool (15-mers overlapping by 11 amino acids; JPT Peptide Technologies, Berlin, Germany); staphylococcal enterotoxin B (5 μg/mL); phorbol 12-myristate 13-acetate (50 ng/mL) with ionomycin (500 ng/mL); or medium containing DMSO (as a negative control; all stimulants were from Sigma except where noted). All peptides were used at 3.5 μg/mL per peptide. Cells were stimulated for 5 hours at 37°C/5% CO_2_ in cell culture medium containing 5 μg/mL brefeldin A (Sigma), 1 μM monensin (BD Biosciences), a FITC-conjugated antibody against the degranulation marker CD107a, and costimulatory antibodies anti-CD28 and anti-CD49d. After stimulation, cells were stained with live/dead fixable aqua dead cell stain and then antibodies recognizing CD4 and CD8. A 4% paraformaldehyde solution was used to fix cells followed by permeabilization with FACS Perm 2 (BD Biosciences). Cells were then stained with antibodies recognizing CD3, interferon-γ, tumor necrosis factor-α, interleukin-2, and macrophage inflammatory protein-1β. Complete antibody identification, including suppliers, catalog numbers, clones, and fluorochromes, is given in **S1 Table**. Cells were acquired within 24 hours on an LSRII flow cytometer (BD Biosciences).

### Drug exposure and measurement

To assess if drugs used in HIV-1 prevention could be affected by the process of cryopreservation and subsequent thawing procedures, colorectal and ectocervical explants were set up as described above for HIV-1 infection. Stocks of dapivirine (non-nucleoside reverse transcriptase inhibitor [NNRTI]; International Partnership for Microbicides, Silver Spring, MD, USA), MK-2048 (integrase strand transfer inhibitor [InSTI], Merck & Co., Kenilworth, NJ USA), and tenofovir (nucleotide reverse transcriptase inhibitor [NRTI]; NIH AIDS Reagent Program, Division of AIDS, NIAID, NIH) were made in DMSO so that at least a 1:100 dilution was made prior to addition to the explants. Dilutions of each drug were applied to the tissues in duplicate and cultured overnight. After washing, one set of the explants was snap frozen to define drug levels in fresh tissues. The other explants were cryopreserved to define drug levels after storage and thawing of viable tissues.

An ultra-performance liquid chromatographic tandem mass spectrometry (UPLC-MS/MS) assay was validated for the determination of MK-2048 in human plasma. The assay range was 25 pg/mL to 50,000 pg/mL with a lower limit of quantitation of 25 pg/mL. The validated method was applied to tissue alternative matrices. Pre-weighed biopsies were digested and extracted with one mL of blank 70% lithium heparin plasma: 30% Collagenase 1a 5.0 mg/mL solution. Samples were vortexed, homogenized, and further diluted with human plasma, up to 10,000-fold, to achieve a concentration within the assay range. Final concentrations were reported as pg/mg, after correcting for dilutions and tissue weight.

Dapivirine concentration was measured on an Acquity UHPLC (Waters Corporation, Milford, MA, USA) with a Quantum Access MAX mass spectrometer (Thermo Fisher). The weighed explants were homogenized in acetonitrile (ACN) and water using a Precellys 24 homogenizer (Bertin Instruments, Rockville, MD, USA). An internal standard of d_4_-dapivirine was added. The samples were treated with 25% NH_4_OH, methyl t-butyl ether, 0.9% NaCl, and then rotated for 30 min. The organic layer was removed and dried. The samples were reconstituted in 50 mM ammonium formate:ACN (40:60 ratio) for analysis. The liquid chromatography conditions used a Hyperclone 3µm BDS C8 150×4.6 mm column (Phenomenex, Torrance, CA, USA) running a gradient of 100% 5mM ammonium formate in 60% ACN to 100% 5mM ammonium formate in 80% ACN over 3.5min. The flow rate was 1mL/min. A positive selective reaction monitoring (SRM) was used with 330.2→158 m/z for dapivirine, and 334→145 m/z for d_4_-dapivirine. Samples were analyzed in the range 0.2 ng/mL to 50 ng/mL. Final concentrations were reported as ng/mg, after correcting for dilutions and tissue weight.

The same Acquity UHPLC was used for tenofovir-diphosphate (TFV-DP) tissue concentration measurement. Tissues were homogenized in 70% ice-cold methanol. After centrifugation, TFV-DP was separated from total TFV using solid phase extraction with Waters QMA cartridges. The supernatant from the homogenization was added and washed thoroughly with 75mM KCl followed by 50mM KCl. TDF-DP was eluted into collection tubes with 1M KCl. The TDF-DP extracts were spiked with a ^13^C_5_-TFV internal standard and dephosphorylated using acid phosphatase. Desalting was accomplished through a second solid phase extraction procedure. The entire dephosphorylated sample was loaded onto Oasis Hydrophilic-Lipophilic Balance (HLB) cartridges (Waters Corporation) and washed with 1% trifluoroacetic acid. TFV-DP (as TFV) was eluted with methanol and concentrated. The samples were then reconstituted in 0.1% formic acid. The LC column was a ZORBAX XDB-C18 5µm, 4.6×50 mm (Agilent Technologies, Santa Clara, CA, USA) maintained at 30°C. Separation was accomplished using a mobile phase of 0.1% formic acid in water and 0.1% formic acid in methanol. The gradient ran at 0.5 mL/min from 5% methanol to 50% methanol over two min and held for 30 sec before re-equilibrating to initial conditions. The mass spectrometer was maintained in positive selective reaction monitoring with 288→176.1 m/z for TFV and 293→181.1 m/z for ^13^C_5_-TFV. Samples were analyzed in the range 0.2 ng/mL to 100 ng/mL. Final concentrations were reported as ng/mg, after correcting for dilutions and tissue weight.

### Data and statistical analysis

For both alamar blue fluorescence and nanoluciferase luminescence, values are background subtracted. For alamar blue, metabolic activity, as measured by fluorescence, was normalized to metabolic activity measured in fresh well. Percent of fresh tissue was determined by dividing the fluorescence measured in individual wells by the average of the wells containing fresh tissues and multiplying by 100.

For colorectal cytokine production and CD107a expression, single-positive gates were placed for each function. The gates were the same for all unstimulated and antigen-stimulated samples. In some cases of SEB and PMA-Ionomycin stimulation, a shift was observed in the negative population of certain functional markers. In these cases, the gates were adjusted on a bivariate plot to better discriminate the boundaries of the negative population. The gates determined on these bivariate plots were used to set the single-positive gates. Once gates were determined, all possible boolean gates were created. Functional responses after stimulation were compared to background responses (DMSO plus costimulatory antibodies) using a previously described statistical test [26], which takes into account the number of gated events rather than percentages. If post-stimulation responses are significantly higher than the background (i.e. p < 0.05), net responses are calculated by subtracting the background; otherwise, responses are set to 0.

We compared preserved samples to fresh samples using a non-inferiority approach. We set a lower effect size bound of 80%, meaning that we regard preserved tissues to be non-inferior to fresh tissues if their assay performance is 80% or greater than that of the fresh tissues. This threshold is similar to quality control thresholds used for thawed peripheral blood mononuclear cells used by the HIV Vaccine Trials Network [27,28]. To test for non-inferiority, we used a two-step procedure. First, we tested for difference between fresh and preserved tissues, with a standard null hypothesis of no difference between fresh and preserved. If that test resulted in a p-value of greater than 0.05, we conducted a second, non-inferiority test. The non-inferiority test used a reversed null hypothesis, namely that the assay performance of preserved tissues was less than 80% that of the fresh tissues. We define a p-value of greater than 0.05 in the test for difference and less than 0.05 in the non-inferiority test as evidence that the preserved tissues are non-inferior. This threshold-based approach is more informative than declaring non-inferiority on the basis of a p-value > 0.05 for the first test only, because that can simply be a consequence of insufficient power. In general, our experiments were weakly powered, with sample sizes in the range of 5-20.

Detailed summary and statistical tables are in **S1 File**. Specific statistical tests are indicated in the text or in the tables. Single comparisons were generally done with paired t-tests and multiple comparisons with repeated measures ANOVA and Tukey post-tests. Survival data was modeled using Cox proportional hazard regression, with non-inferiority concluded if the 95% confidence interval around the hazard ratio excluded the 80% threshold. Data analysis was done using R version 3.4.1 [29] and the packages Hmisc [30], broom [31], survival [32], tidyverse [33], stringr [34], forcats [35], lazyeval [36], readxl [37], plater [38], and nlme [39]. The package ggplot2 [40] was used for figures and pander [41] and knitr [42] were used for tables. Analysis code and experimental data are included in S2 File.

## Results

We set out to design optimal cryopreservation and vitrification protocols for colorectal and cervicovaginal tissues. We began with a series of preliminary experiments using only vaginal tissues. For cryopreservation, we placed individual 2 mm explants (pieces of tissue dissected in the laboratory from surgically excised vaginal epithelium) in cryovials containing 200 µL 10% DMSO in fetal bovine serum and froze the tissues at 1°C/minute. Our vitrification medium was 20% DMSO with 20% EG in a solution of phosphate-buffered saline containing 20% fetal bovine serum.

### Optimization of vitrification

We started by testing different methods of vitrification. For vitrification to occur the cooling must be as rapid as possible and the tissues must be permeated with the vitrification medium. To permeate the tissues, we soaked them in a half concentration solution of vitrification medium (10% EG and 10% DMSO) for 10 min, and then in a full concentration solution (20% and 20%) for 10 more min, both at room temperature. For the freezing portion, we tested the presence/absence of excess vitrification medium (“wet” vs. “dry”) and the best container for holding tissue pieces. We therefore put the explants into cryovials, into vitrification straws, or onto pieces of aluminum foil. The explants were either blotted dry and put into the containers (“dry”) or put into the containers along with 200 µL vitrification medium (“wet”). In each case, the container and explant were plunged directly into liquid nitrogen and held there until bubbling subsided.

To get an indication of tissue viability, we measured cellular metabolic activity. The thawed or fresh explants were cultured for 5 hours in the presence of alamar blue (resazurin), which is reduced to the fluorescent resorufin by the chemical reducing potential created by metabolic activity of live cells. As indicated in **Fig 1A**, higher levels of fluorescence were seen in the “dry” conditions than in the “wet” conditions, suggesting that blotting the tissues dry is better than freezing in the vitrification medium. Additionally, the straw and foil seemed to be better than the cryovial, perhaps due to the thickness of the cryovial plastic slowing the freezing rate.

**Fig 1.**
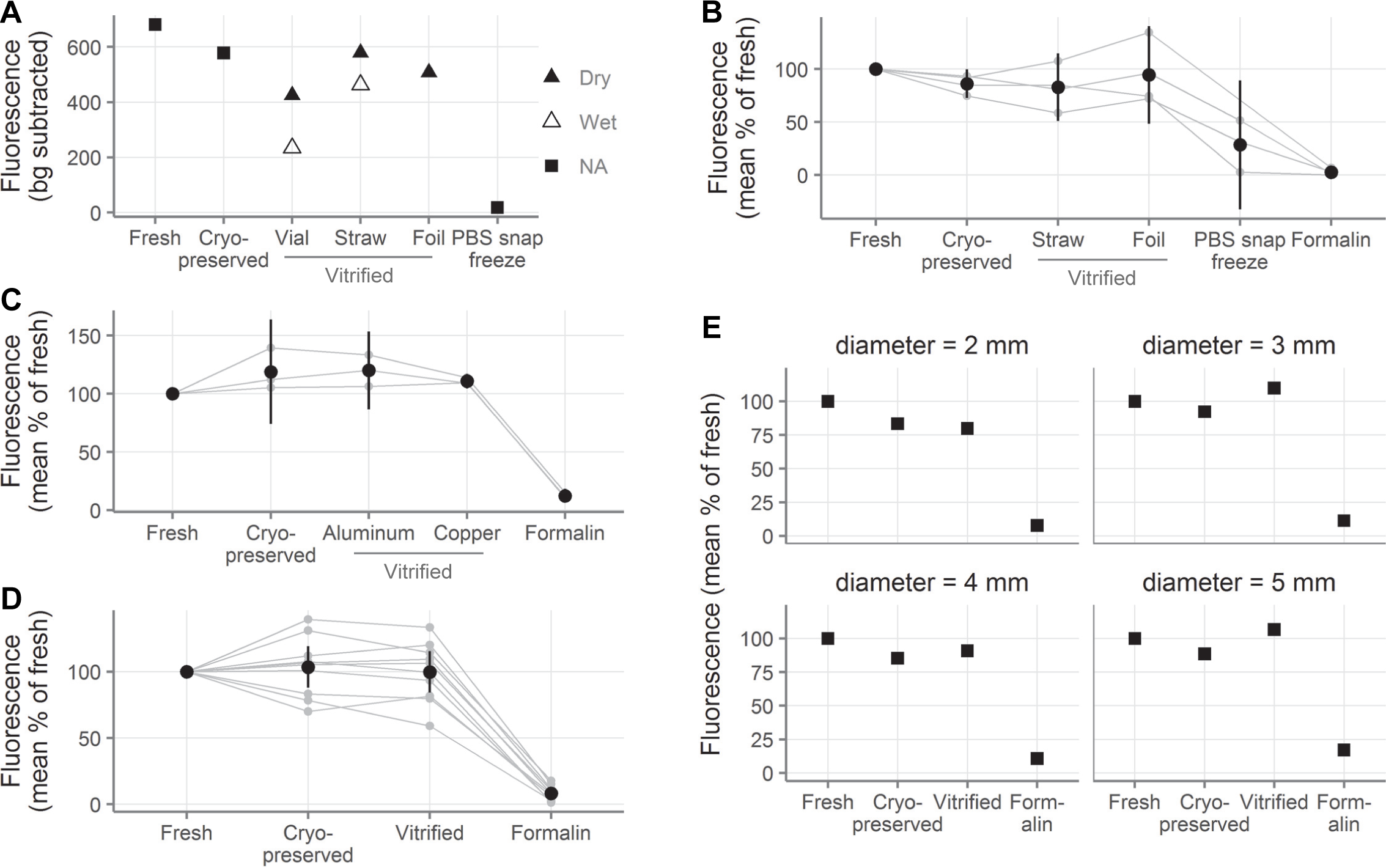
Cryopreserved and vitrified vaginal explants show similar levels of metabolic activity to fresh explants. **A**, Background-subtracted alamar blue fluorescence of vaginal explants after 5h of culture following cryopreservation and vitrification. “Wet” explants were immersed in vitrification medium during freezing and “dry” explants were blotted dry before freezing (n = 1 tissue donor). “NA” stands for “not applicable” and indicates non-vitrification conditions; “PBS”, “phosphate buffered saline”; “bg”, “background.” **B**, Normalized alamar blue fluorescence of vaginal explants (expressed as percent of the fluorescence measured in the fresh explants from the same donor). Promising conditions from A were repeated with explants from 3 tissue donors, which are shown together with the data from the first tissue donor in A. **C**, Normalized alamar blue fluorescence of vaginal explants comparing vitrification with aluminum or copper foil (n = 3 tissue donors). **D**, Summary of experiments comparing cell metabolic activity after cryopreservation and vitrification (aluminum foil) in vaginal explants (n = 10 tissue donors). **E**, Effect of tissue size on cell metabolic activity after cryopreservation or vitrification (aluminum foil). Here fluorescence was normalized to the fresh explants of the same diameter (n = 1 tissue donor; “mm”, millimeter). In A and E, symbols indicate the average of two to three explants. In B-D, smaller gray points indicate the average of duplicate explants, with gray lines indicating samples from the same tissue donor. Black points show the mean across all tissue donors and black vertical lines show the 95% confidence interval of the mean.

Based on this experiment, we repeated the most promising conditions with tissues from an additional three donors. As shown in **Fig 1B**, the cryopreserved and vitrified explants were nearly as metabolically active as the fresh tissue. In contrast, snap freezing the tissue in PBS or fixing it with formalin overnight almost completely eliminated metabolic activity. Based on the apparent equivalence of the dry straw and foil, we used foil for further experiments because it is more convenient.

Given that we chose to proceed with the foil method, we wanted to know whether the foil material would have an effect. In particular, the cooling rate might be enhanced by a material that conducts heat more quickly. We therefore compared aluminum foil, with a thermal conductivity of 205 watts per meter kelvin (W/(m K)), to copper, with a thermal conductivity of 401 W/(m K). **Fig 1C** shows that the type of foil used had no noticeable effect on the success of the vitrification in experiments with explants from three donors.

### Summary of all tissue viability assessments

In total, we conducted experiments with samples from 10 tissue donors comparing cryopreservation and foil vitrification by the alamar blue assay. As shown in **Fig 1D** and **Table A in S1 File**, explants preserved with foil vitrification or cryopreservation had similar levels of cellular metabolic activity to fresh controls. Specifically, cryopreserved explants had 103.5% (95% CI: 87.8-119.2) as much activity as fresh explants, while vitrified explants had 99.8% (84.1-115.4). To determine whether the preserved tissues were non-inferior to the fresh tissue, we used the non-inferiority approach described in the methods. We defined non-inferior as a maximum of 20% below the fresh tissues, as shown in **Table B in S1 File**. Neither preservation technique differed significantly from fresh by paired t-tests for difference (null hypothesis: no difference between metabolic activity of fresh and preserved, all p > 0.05). In the second stage of non-inferiority testing, we conducted paired, one-sided t-tests (null hypothesis: preserved metabolic activity less than 80% of fresh). Both preservation techniques were non-inferior to fresh at our quality control threshold of 20% (all p < 0.05).

### Effect of tissue size on preservation

Now that we had a promising set of protocols, we wanted to assess whether the protocols would work for tissue pieces of different sizes. All of the previous experiments were conducted with vaginal explants obtained by 2 mm punch biopsy (cylindrical explants 2 mm in diameter and 2-4 mm thick). Here, we froze tissues obtained with 2, 3, 4, and 5 mm biopsy punches (volume from 6 to 80 cubic mm). **Fig 1E** shows the results of this experiment with tissue from one donor. The results suggest that for explants up to at least 5 mm in diameter, both cryopreservation and vitrification seem to work equally well in preserving metabolic activity.

### Vaginal tissue structure after preservation

To assess how cryopreservation and vitrification affect tissue morphology, we performed hematoxylin and eosin (H&E) stains of sectioned vaginal tissue. We took three explants each from five donors and embedded and stained them either fresh or after cryopreservation or vitrification.

Representative images from a single donor are shown in **Fig 2**. All slides were reviewed by a pathologist, who was blinded to treatment, and determined that no gross changes in tissue structure were apparent in samples from any of the five donors. The only noticeable difference was that red blood cells could be seen in tissues that were fixed fresh (visible in the fresh tissue in **Fig 2**), but not in the preserved tissues.

**Fig 2.**
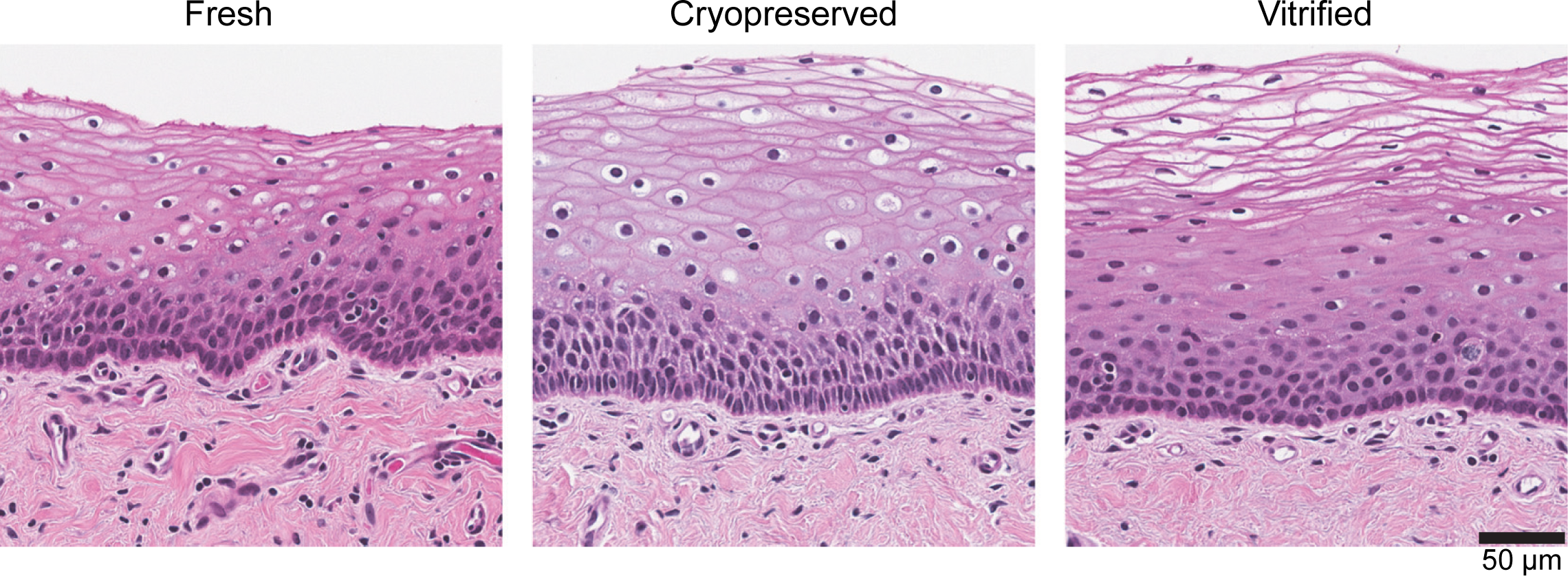
Cryopreserved and vitrified vaginal explants maintain similar tissue structure to fresh explants. Example images from one tissue donor of hematoxylin-and-eosin-stained sections of vaginal tissue after no treatment (left), cryopreservation (middle), and vitrification (right) (total n = 5 tissue donors). Scale bar indicates size (“µm”, micrometer).

### HIV-1 infection of vaginal tissues

We next challenged fresh and preserved explants with HIV-1 to determine whether the tissue would remain equally susceptible to infection after preservation.

We used a nanoluciferase-secreting HIV construct to monitor infection over time [19]. After thawing, we challenged 2 mm explants with HIV, in parallel with fresh biopsies, as described in the Methods and maintained them for three weeks with regular supernatant collection.

**Fig 3A** shows an example infection with vaginal tissue from one donor, where three explants per condition were challenged with virus. In this experiment, 2/3 fresh explants, 3/3 cryopreserved, 2/3 vitrified, and 0/3 formalin-fixed explants became infected.

**Fig 3.**
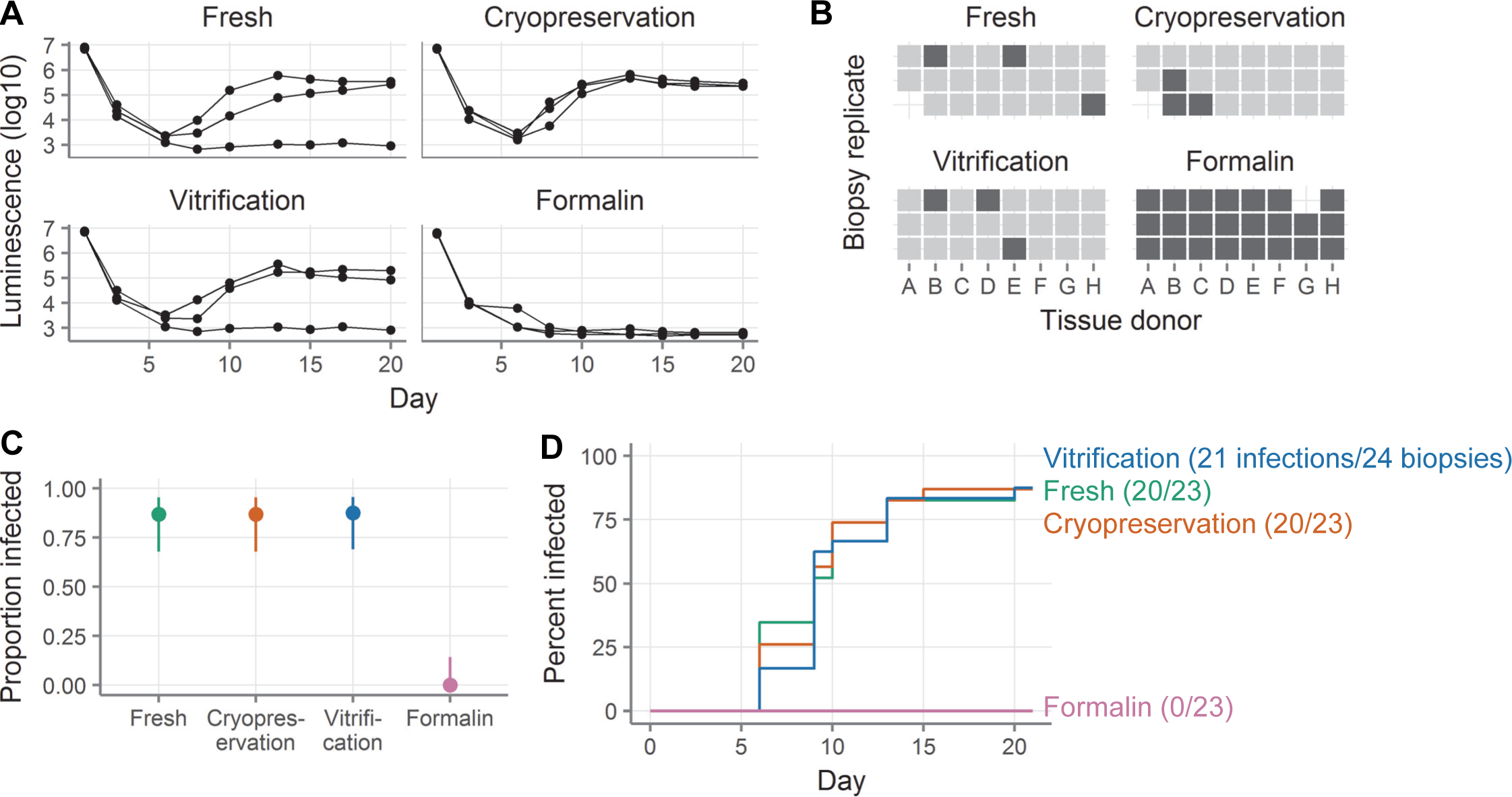
Cryopreserved and vitrified vaginal explants are similarly susceptible to HIV infection as fresh explants. **A**, HIV challenge curves showing nanoluciferase luminescence from a single tissue donor. Each line represents a single explant and points represent supernatant collections. The left-most point shows the viral inoculum and the next point shows the level of residual nanoluciferase after virus wash-off. All three explants come from the same donor. **B**, Summary of HIV challenge experiments. Explants are defined as infected on the first day after day 5 where the luminescence exceeds 1E4. Each square represents a single explant, with each column representing a different donor (total n = 8 donors with 2-3 explants per donor per condition). Light squares indicate explants that became infected and dark squares explants that did not. Missing squares indicate cases where there was inadequate tissue available to have three explants in all conditions or where an explant was lost by experimental error. **C**, Proportion of explants that became infected. Points show the proportion of all challenged explants that became infected. Lines show the 95% binomial proportion confidence intervals. **D**, Survival curves indicating cumulative HIV infection for all explants tested in each condition. Fresh explants are shown in green, cryopreserved in orange, vitrified in blue, and formalin-fixed in pink.

We repeated this experiment with explants from 8 donors. As shown in **Fig 3B** and summarized in **Fig 3C**, 20/23 fresh explants, 20/23 cryopreserved, 21/24 vitrified, and 0/23 formalin-fixed explants became infected. The day of infection was defined as the first day after day 5 on which luminescence exceeded 10,000 units. The fresh, cryopreserved, and vitrified explants all got infected at similar rates (hazard ratio relative to fresh for cryopreservation 1.03, for vitrification 0.95, all p > 0.05, see **Fig 3D** and **Table C in S1 File**). However, the confidence intervals for the hazard ratios were too wide to exclude our quality control threshold of 0.8, so we cannot formally conclude non-inferiority of HIV infectability.

### Cell yields from cryopreserved and vitrified colorectal biopsies

We next wanted to know how cryopreservation and vitrification compared in colorectal tissue. We obtained 30 colorectal biopsies (pieces of tissue obtained from donors through flexible sigmoidoscopy) from each of five donors. Fifteen biopsies were cryopreserved and 15 were vitrified per donor. After thawing, biopsies were digested to obtain a single-cell suspension and leukocytes were isolated by Percoll density gradient centrifugation. The yield of total cells was determined by Guava Viacount assay and then cell phenotyping was performed, with gating done as shown in **S1 Fig**.

As shown in **Fig 4A**, cryopreserved colorectal biopsies yielded dramatically more cells than vitrified biopsies. The mean total yield from cryopreserved biopsies was 6.84 million (95% CI: 4.44-9.21 million) and only 0.27 million (−0.02-0.56 million) from vitrified biopsies (**Tables D-E in S1 File**, p = 0.0018 by paired t-test). In other words, the yield from vitrification was less than 10% of the yield from cryopreservation.

**Fig 4.**
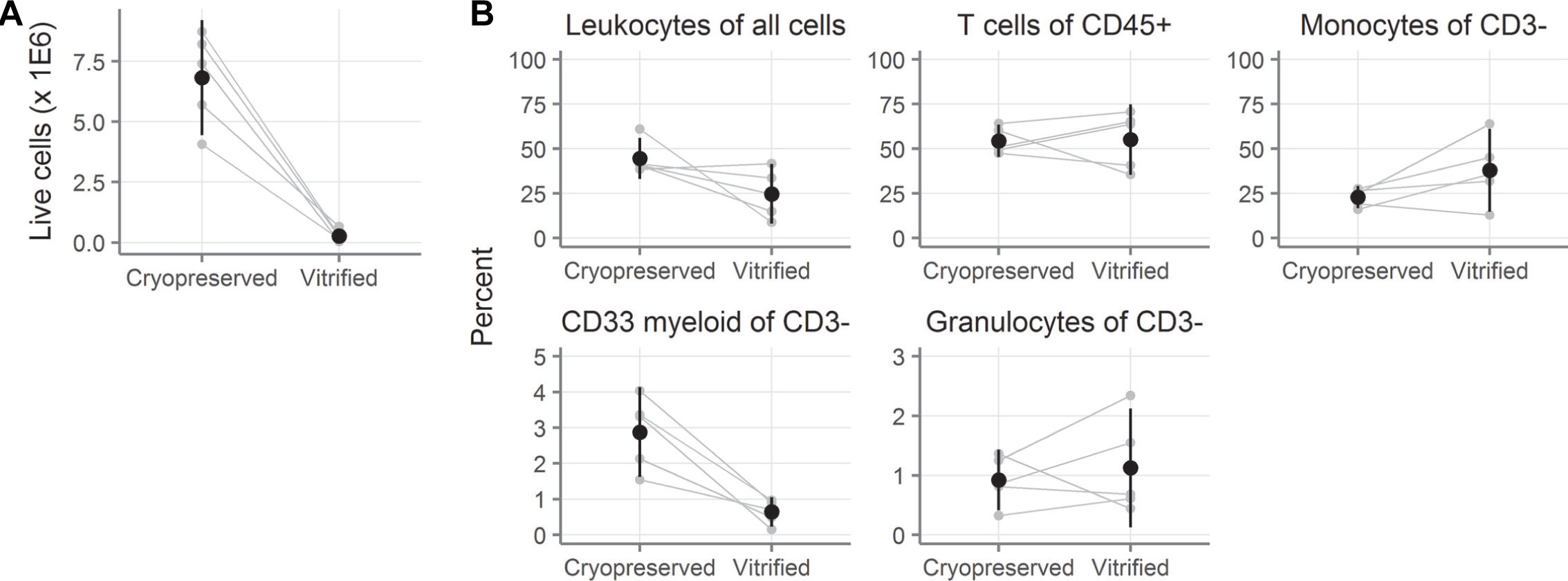
Ten-fold more cells are recovered after cryopreservation than vitrification of colorectal biopsies. **A**, Number of live cells recovered after digestion of colorectal biopsies that had been cryopreserved or vitrified (n = 5 donors with 15 biopsies per donor per condition). **B**, Cell phenotypes as measured by flow cytometry. Leukocytes are defined as CD45^+^. T cells are CD45^+^CD3^+^. Monocytes are CD45^+^CD3^−^CD13^+^. CD33 myeloid are CD45^+^CD3^−^CD33^+^. Granulocytes are CD45^+^CD3^−^CD66b^+^. Small gray symbols indicate the number (A) or percent (B) from fifteen colorectal biopsies, with gray lines indicating samples from the same tissue donor. Black symbols show the mean across all tissue donors and black vertical lines show the 95% confidence interval of the mean.

There were a few differences between the two conditions in the phenotypes of the cells that were recovered (**Fig 4B** and **Tables F-G in S1 File**). The main differences were that fewer of the viable cells were CD45^+^ in vitrified biopsies (24.7% [8.0-41.4]) than in cryopreserved (44.5% [33.0-56.0]; p = 0.10) and that vitrification reduced the frequency of CD33+ cells more than four-fold. A complete summary is available in **Tables F-G** in **S1 File**. To conclude, dramatically more cells were recovered from cryopreserved colorectal biopsies than vitrified ones.

### HIV infection of colorectal and ectocervical tissues

After determining that cryopreservation was markedly superior to vitrification for colorectal leukocyte recovery, we decided to stop testing vitrification. We went on to further validate cryopreservation in additional tissue types and assays. We therefore repeated our HIV infection experiments using both colorectal and ectocervical explants, comparing cryopreserved to fresh tissues.

**Fig 5A-D** show HIV challenges of colorectal explants. We evaluated colorectal explants from 12 donors and infection curves from one representative donor are shown in **Fig 5A**. Explants are defined as infected on the first day after day 5 where the p24 concentration exceeds 250 pg/mL. As summarized in **Fig 5B-C**, we saw infection in 27/30 fresh explants and 23/31 explants cryopreserved in 10% DMSO. Based on our prior work [1], we also tested cryopreservation with 6% DMSO, 5% EG and 50 mM trehalose, where 10/18 explants were infected. Infection occurred at a lower rate in cryopreserved than in fresh explants, particularly those cryopreserved with 6% DMSO, 5% EG and 50 mM trehalose (hazard ratio relative to fresh 0.42 [95% CI 0.2-0.86], p = 0.02; **Fig 5D** and **Table H in S1 File**). While the hazard ratio for explants cryopreserved with 10% DMSO was below 1 (0.7 [0.4-1.23), we could neither conclude that the infection rate was lower than that of fresh (p > 0.05), nor that it was non-inferior, because the lower bound of the 95% CI was less than 0.8. For most donors, the rates of infection between fresh and 10% DMSO were similar, but in three donors (donors D, E, and F), the rate was much lower in the cryopreserved explants.

**Fig 5.**
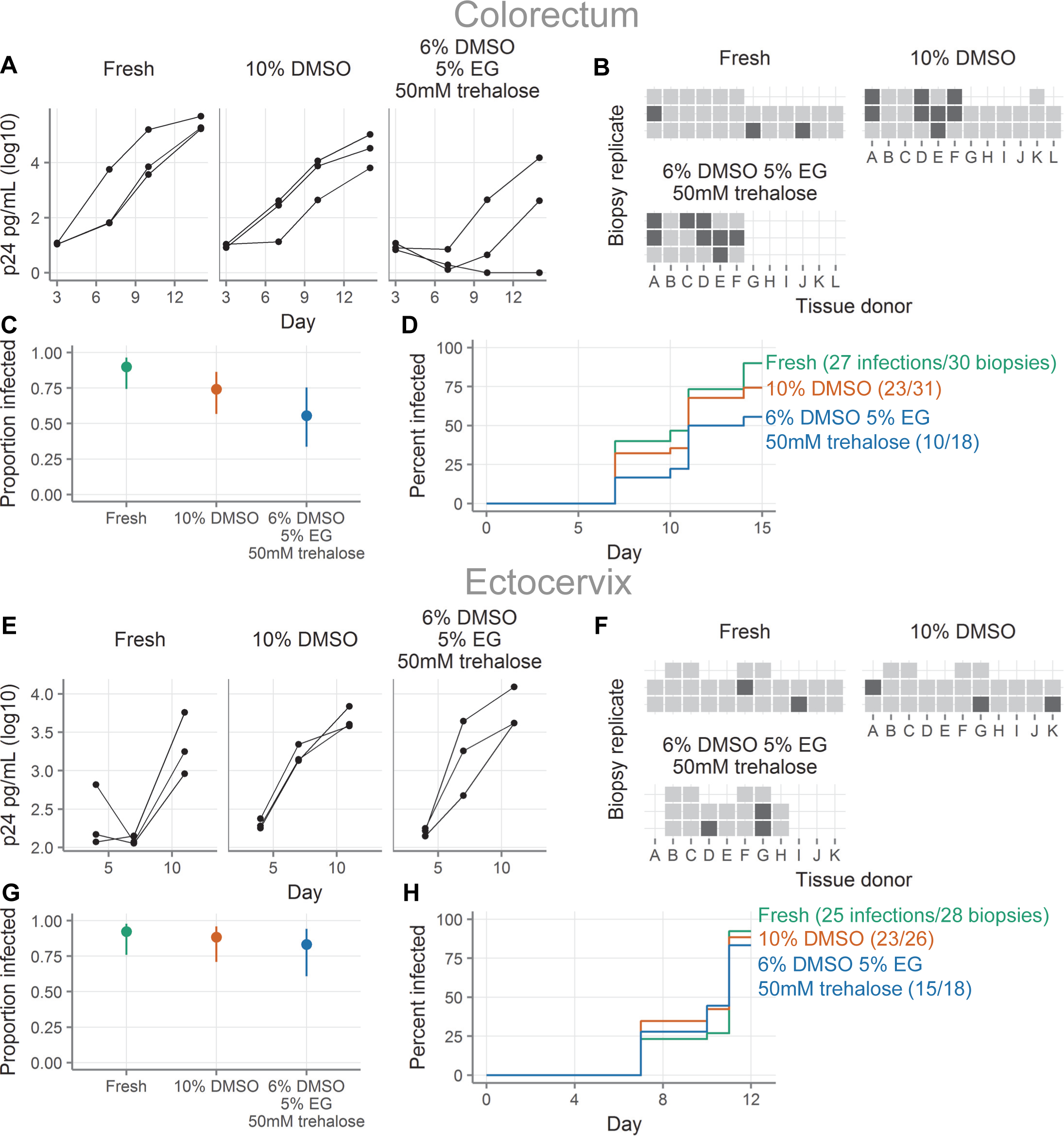
Cryopreserved colorectal explants are somewhat less susceptible to HIV infection than fresh colorectal explants, while cryopreserved and fresh ectocervical explants are similarly susceptible. **A**, HIV challenge curves for colorectal tissue showing HIV p24 concentrations (picograms/milliliter) from a single tissue donor either fresh or after cryopreservation with the indicated cryopreservation media. Each line represents a single explant and points represent supernatant collections. All three explants come from the same donor. “DMSO”, dimethylsulfoxide; “EG”, “ethylene glycol”; “mM”, “millimolar”. **B**, Summary of HIV challenge experiments for colorectal tissues. Each square represents a single explant, with each column representing a different donor (total n = 12 donors with 2-3 explants per donor per condition, as shown). Light squares indicate explants that became infected and dark squares explants that did not. Missing squares indicate cases where there was inadequate tissue available to have three explants in all conditions or where an explant was lost by experimental error. Explants are defined as infected on the first day after day 5 where the p24 concentration exceeds 250 pg/mL. **C**, Proportion of colorectal explants that became infected. Points show the proportion of all challenged explants that became infected. Lines show the 95% binomial proportion confidence intervals. **D**, Survival curves indicating cumulative HIV infection for all colorectal explants tested in each condition. Fresh explants are shown in green, cryopreserved with 10% DMSO in orange, and cryopreserved with 6% DMSO, 5% EG, and 50 mM trehalose in blue. **E**, HIV challenge curves for ectocervical tissue, as in A. **F**, Summary of HIV challenge experiments for ectocervical tissue, as in B, with a threshold of infection of 500 pg/mL (total n = 11 donors with 2-3 explants per donor per condition, as shown). **G**, Proportion of ectocervical explants that became infected, as in C. **H**, Survival curves for HIV infection of ectocervical explants, as in D.

**Fig 5E-H** show HIV challenges of ectocervical explants. Infection curves from a representative donor are shown in **Fig 5E**. Explants are defined as infected on the first day after day 5 where the p24 concentration exceeds 500 pg/mL (this threshold is higher than the threshold for colorectal explants because ectocervical explants have higher background levels of p24, as can be seen by comparing **Fig 5A** and **Fig 5E**). **Fig 5F-G** summarize infections with explants from 11 donors. We saw infection in 25/28 fresh explants, 23/26 explants cryopreserved in 10% DMSO, and 15/18 explants cryopreserved in 6% DMSO, 5% EG and 50 mM trehalose. The rates at which the explants became infected were similar for fresh and cryopreserved (**Fig 5H** and **Table I in S1 File**). However, the confidence intervals for the hazard ratios were too wide to exclude our threshold of difference of 0.8, so we cannot formally conclude non-inferiority.

In conclusion, as with vaginal tissues, fresh and cryopreserved ectocervical explants were infected equally well. In colorectal explants, a smaller proportion of cryopreserved explants became infected than fresh explants.

### Comparison of cryopreserving whole biopsies and isolated cell suspensions

We next sought to determine whether it is better to (1) isolate cells from a biopsy and cryopreserve the cell suspension or (2) cryopreserve the biopsy and isolate the cells after thawing. In these experiments, we expanded a previously published experiment ([1], **Fig 6B-C**). In the previous study, we reported on CD8 T cell responses after cryopreservation from five HIV^−^ donors. Here, we enlarge that dataset to include nine HIV^+^ and nine HIV^−^ individuals, who each donated 30 colorectal biopsies. Ten biopsies were cryopreserved intact. The remaining 20 were digested to isolate cells. Half of the cells were cryopreserved and the other half were immediately assayed for cytokine production. After 1-5 months, the cryopreserved biopsies and cell suspensions were thawed. The biopsies were digested to cell suspensions and both sets of cell suspensions were assayed for cytokine production.

**Fig 6.**
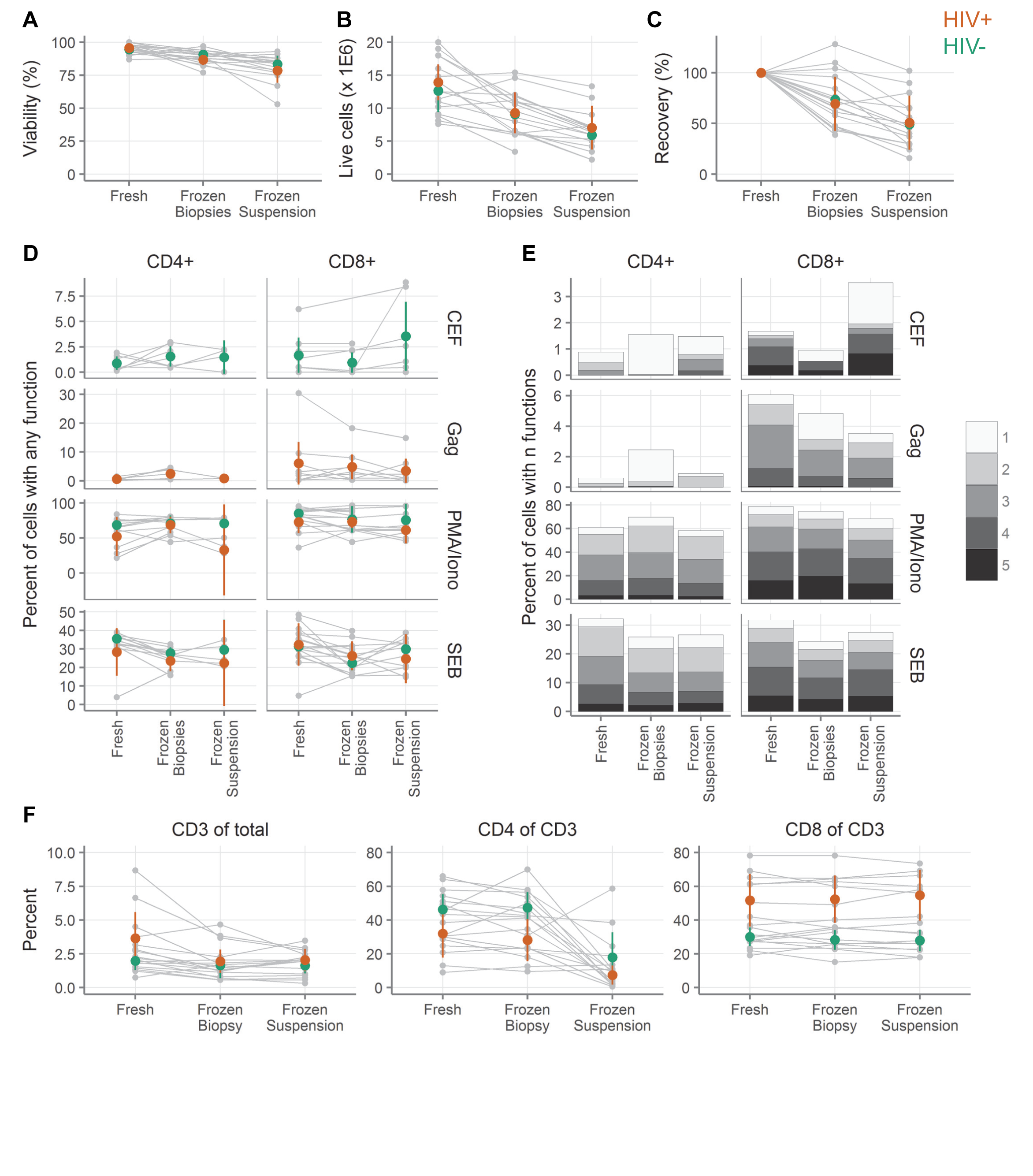
Cryopreservation of intact colorectal tissue leads to greater cell numbers than cryopreservation of cell suspensions, while cellular functionality is similar. **A**, Viability by trypan blue exclusion of colorectal cells digested from biopsies either fresh, cryopreserved as biopsies, or cryopreserved after digestion (n = 18 tissue donors). **B**, Live cell yield by trypan blue exclusion. **C**, Recovery of live cells after cryopreservation relative to fresh samples from the same donor. **D**, Background-subtracted cytokine production from colorectal T cells after stimulation with cytomegalovirus, Epstein-Barr virus and influenza virus peptides (CEF); HIV Gag peptides (Gag); phorbol 12-myristate 13-acetate and ionomycin (PMA/Iono); or staphylococcal enterotoxin B (SEB). Small gray symbols indicate individual samples, with gray lines connecting samples from the same tissue donors. Colored symbols show the mean across all samples from HIV^+^ (orange) and HIV^−^ (green) donors, with vertical lines showing the 95% confidence interval of the mean. **E**, Polyfunctionality of colorectal responses to stimulation. Each sub-bar corresponds to the percent of cells with the indicated number of functions, averaged across the donors. Total bar height indicates the total percent of cells with one or more functions.

Total cell number and viability were determined by trypan blue exclusion. Fewer cells were recovered with both methods of cryopreservation than with fresh samples, but cryopreservation of biopsies was considerably better than cryopreservation of cell suspensions (**Fig 6A-C**, **Tables J-S in S1 File**). The viability was 95.1% (95% CI: 93.2-96.9; n = 9 HIV^+^, 9 HIV^−^) in the fresh samples, 88.7% (86.2-91.2; n = 9 HIV^+^, 9 HIV^−^) in the frozen biopsies, and 81.1% (76.2-86.1; n = 8 HIV^+^, 9 HIV^−^) in the frozen suspensions (**Fig 6A**, **Tables J-M in S1 File**, p < 0.05 for all pairwise comparisons by ANOVA with Tukey post-tests). The sample size is one lower (17 instead of 18) for frozen suspensions because in one case insufficient biopsies were available. In total, we recovered an average of 13.3 million live cells (11.4-15.2 million) from the fresh samples, 9.2 million (7.6-10.8) from the frozen biopsies, and 6.5 million (5.0-8.0) from the frozen suspensions (p < 0.05 for all pairwise comparisons by ANOVA with Tukey post-tests, except the comparison of frozen suspensions to frozen biopsies where p = 0.05). Given that the differences in cell number were considerably larger than 20%, none of the samples meet our definition of non-inferiority. The average recovery of viable cells relative to the fresh sample was 71.4% (58.9-83.8) for frozen biopsies and 49.8% (37.5-62.1) for frozen suspensions (**Fig 6C**, **Tables R-S in S1 File**). Means and 95% confidence intervals are shown separately for HIV^+^ and HIV^−^ donors, but were not compared statistically.

While freezing biopsies had a clear advantage over freezing cell suspensions in terms of cell yield and viability, we also wanted to determine whether cytokine production and other cellular functions were affected by freezing condition. For these experiments, in many cases we could compare fresh, frozen biopsy, and frozen suspension, but in some cases the cell yield was too low to include a particular test. In the text and figures, we show all of the samples and tests where we had data for the fresh condition and at least one of the frozen conditions. We additionally calculated the statistics on only those samples and tests where we could compare all three conditions (“fully paired”). The statistics for both the fully paired and complete data sets are displayed in the tables in **S1 File**.

To test T cell function, we stimulated the cells with the mitogen PMA/ionomycin; the superantigen staphylococcal enterotoxin B (SEB); peptides from the commonly encountered pathogens cytomegalovirus, Epstein-Barr virus and influenza virus (CEF) (HIV^−^ donors only); or HIV Gag peptides (HIV^+^ donors only). After stimulation, we measured T cell production of CD107a, representing cell contact-dependent cytotoxic effector function [43], and four cytokines, representing paracrine immune effector function: interferon-γ, interleukin-2, macrophage inflammatory protein (MIP)-1β, and tumor necrosis factor-α, with gating performed as shown in **S2 Fig**. **Fig 6D** shows the percent of cells expressing at least one of those cytokines after stimulation (**Tables T-U** in S1 File). The confidence intervals are largely overlapping with no obvious and consistent differences between the conditions.

Though there seemed to be little difference between fresh and frozen when considering the production of any cytokine, we still wondered whether the polyfunctionality of the responses might be affected. We therefore assessed how many cytokines or CD107a were produced per cell. As shown in **Fig 6E**, the patterns were quite similar regardless of whether the samples were processed fresh, frozen as a cell suspension, or frozen as biopsies.

Finally, we compared T cell frequencies after stimulation (**Fig 6F**, **Tables V-AC** in S1 File). There were somewhat more live T cells as a fraction of all events in fresh samples (2.82% [1.81-3.82]), than cryopreserved biopsies (1.80% [1.23-2.38], p = 0.10) or cryopreserved cell suspensions (1.82% [1.37-2.27], p = 0.11), but there was no difference between the cryopreserved conditions. There was no difference between any of the conditions in the fraction of T cells that expressed CD8, but there was a dramatic reduction in the fraction of T cells that expressed CD4 in the frozen suspension compared to either fresh or frozen biopsies (from nearly 40% to 13%, p < 1.3E-4, **Table Z** in S1 File).

In conclusion, we take the results of these experiments as an indication that it is preferable to cryopreserve intact biopsies rather than cell suspensions. In both cases, the cell yield and viability will be reduced relative to a fresh sample, but less so for the cryopreserved biopsies. Both methods of preservation maintain the functionality of the cells in a state similar to the fresh state.

### Microbicide drug concentrations in cryopreserved tissue

In microbicide trials, it is of interest to measure the amount of drug in tissue after product use by a participant. Samples for this purpose are typically frozen dry. If it were possible to use cryopreserved samples to measure drug concentrations, it would afford investigators more flexibility in sample use. To assess the impact of cryopreservation on tissue drug concentration, we exposed fresh ectocervical and colorectal tissues to tenofovir, dapivirine, or MK-2048 overnight. For each drug, parallel explants were exposed to different concentrations of drug. After exposure, we either snap froze the biopsies immediately or cryopreserved and thawed them before snap freezing for drug concentration measurements. Two explants were used from each donor for each drug, concentration, and treatment (fresh or cryopreserved). The concentrations from these two explants were then averaged. Averaged concentrations from three donors per tissue type are shown in **Fig 7A**. Cryopreserved drug concentrations as a percentage of the fresh concentrations are shown in **Fig 7B**. Complete details are shown in **Table AD in S1 File**. In all cases but one, drug concentrations were lower after cryopreservation compared with fresh samples. Tenofovir diphosphate concentrations, which were only measured in ectocervical explants, tended to be much lower in cryopreserved than fresh explants (3% at the lower exposure level, 37% at the higher exposure level). The reported concentrations for tenofovir likely overestimate the recoveries as several values for cryopreserved explants fell below the lower limit of detection and were set to the lower limit of detection. Concentrations also tended to be lower for dapivirine and MK-2048 after cryopreservation, but to a smaller extent. For MK-2048, concentrations after cryopreservation were reduced similarly between ectocervical and colorectal tissues, with concentrations of 42-97% of fresh. A similar pattern was seen with dapivirine, with cryopreserved drug levels from 44-96%, except in colorectal tissue at the lowest exposure dose, where the mean concentration in cryopreserved tissues was 664% of that of the fresh tissues; it should be noted that the absolute levels measured in this condition were very low. With data from only three tissue donors per condition, the conclusions drawn from these experiments are limited, but the data suggest that the cryopreservation procedure reduces drug concentrations relative to fresh samples, in particular for tenofovir diphosphate.

**Fig 7.**
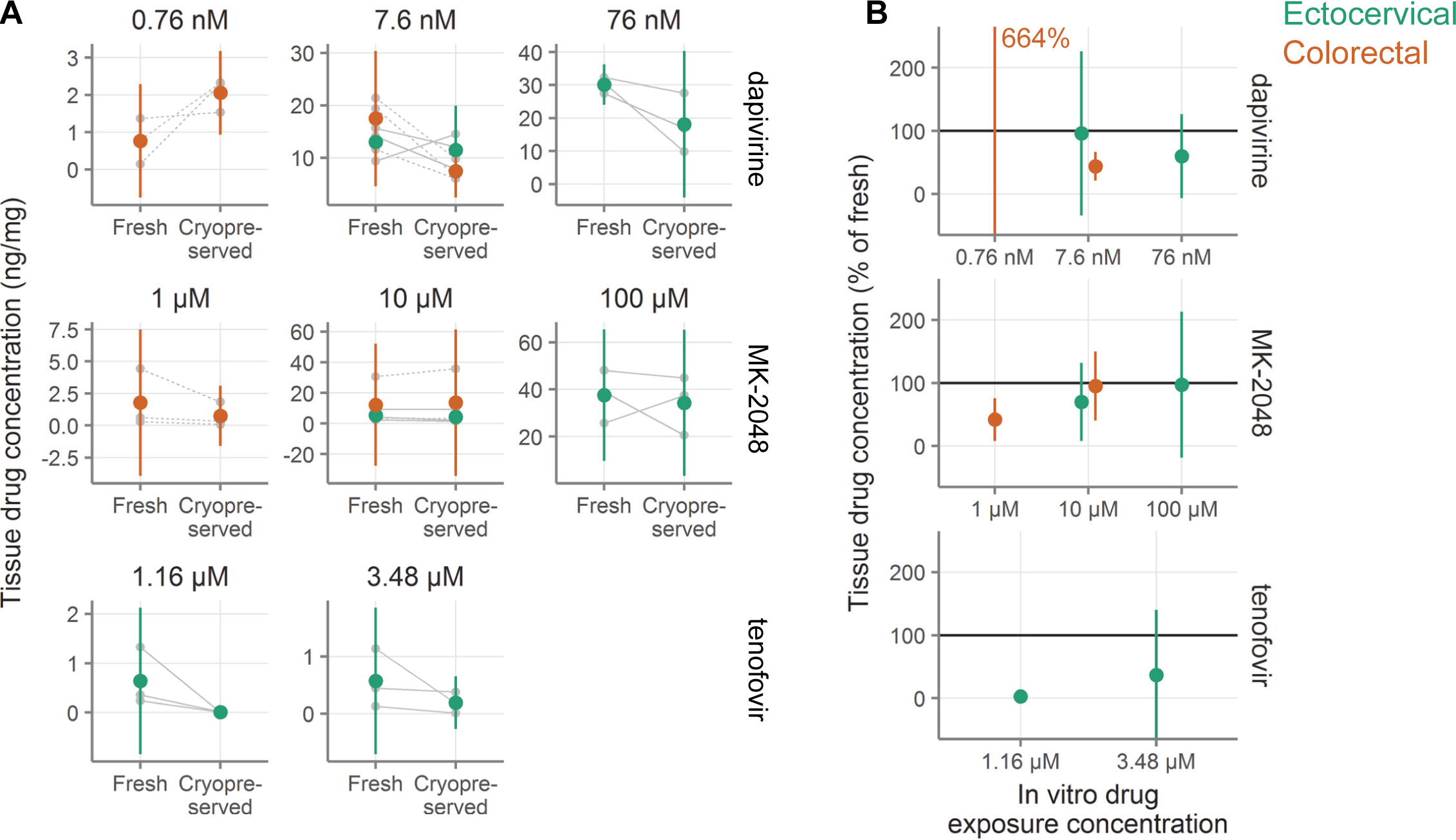
Retention of antiretroviral drugs in ectocervical and colorectal tissue after cryopreservation. **A**, Concentration of drug measured in tissues either fresh or after cryopreservation (n = 3 tissue donors per condition with two explants per donor per condition). Explants were exposed in vitro to antiretroviral drugs at the concentrations indicated above each plot. After drug exposure and cryopreservation, drug levels were measured in the tissues. Small gray symbols indicate the average of two explants and gray lines connect samples from the same tissue donor (dashed indicates colorectal, solid indicates ectocervical). Large symbols indicate the mean for that tissue type (orange indicates colorectal, green indicates ectocervical). Vertical green and orange lines show 95% confidence intervals. **B**, Normalized drug concentration expressed as percent of drug concentration in fresh samples from the same tissue donor. Colored symbols show the mean across all tissue donors and colored vertical lines show the 95% confidence interval of the mean. The mean recovery for dapivirine in colorectal tissue is off-scale and the value is indicated in text.

## Discussion

We conclude that cryopreservation of intact cervicovaginal and colorectal tissues with DMSO works well in a range of assays (**Fig 8)**, while the utility of vitrification is more limited. Cell yields are better with cryopreservation of tissue pieces (about 70% of fresh) than with cryopreservation of isolated cells (about 50%), but other functions are similar. We therefore recommend, when possible, cryopreservation of intact tissue. In **S1 Text** and at https://dx.doi.org/10.17504/protocols.io.p5adq2e, we provide detailed protocols for cryopreservation and vitrification of mucosal tissues. These protocols, in complement with our previously published protocols for mucosal cell suspensions [1], should allow investigators to conduct functional studies with cervicovaginal and colorectal specimens. An additional benefit of cryopreservation of intact tissue is that little sample processing is needed, thus making it feasible to obtain samples at clinical sites where limited laboratory capacity is available. Cryopreservation of samples gives the investigator more flexibility and assay options in the future. Instead of pre-determining the assay to perform and storing the specimen accordingly, samples can be cryopreserved, retaining their viability and structure. At a later date, the viable samples can be thawed and processed according to the needs of the assay.

**Fig 8.**
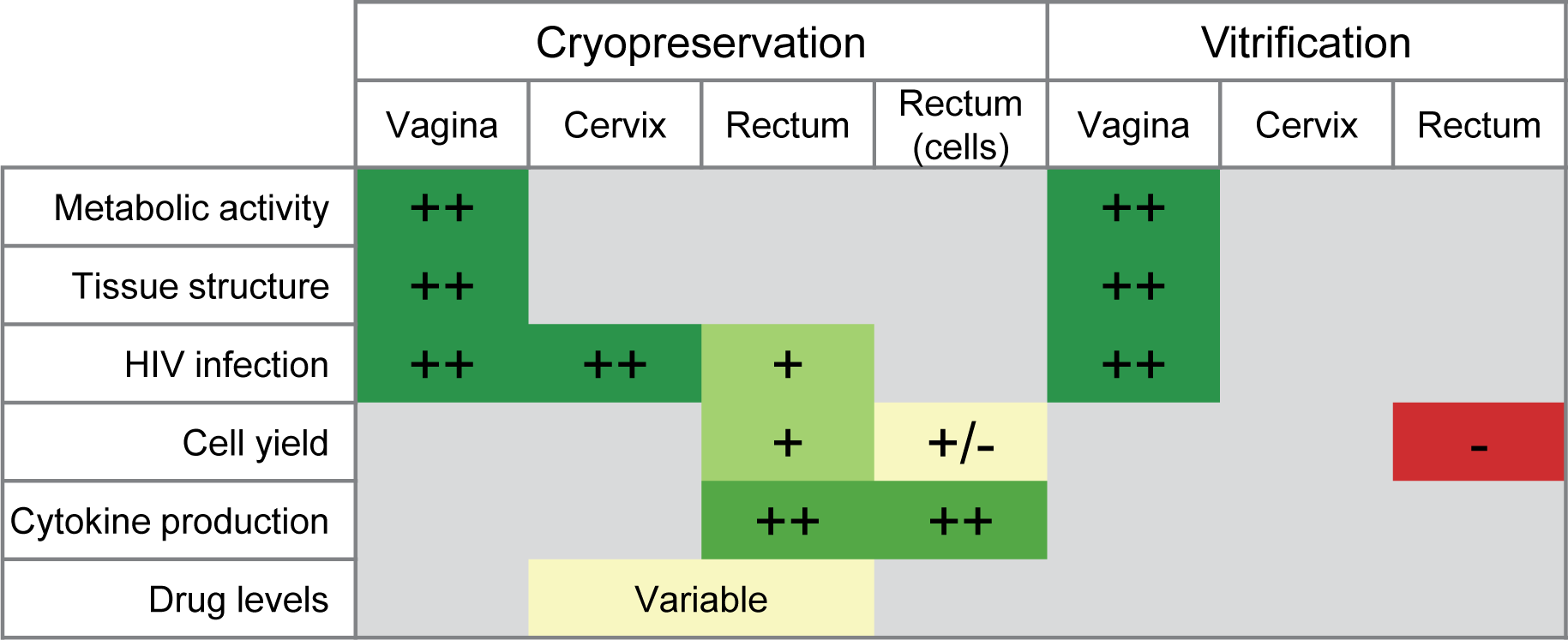
Summary of all cryopreservation and vitrification experiments and tissue types. Signs indicate performance of each tissue type in each assay, relative to the performance of the fresh tissue. ++ indicates equivalence to fresh, + indicates a reduction but probably acceptable,+/− indicates marginal performance, and - indicates inadequate. Colors also indicate performance, with red to yellow to green representing worse to better performance, as compared to fresh tissue. Gray squares indicate that the assay was not performed for that combination of sample type and preservation type.

A limitation of our study is that, as depicted in Fig 8, we were not able to perform each assay in each type of tissue. Therefore, while we can say that cryopreservation works well in a range of tissue types and in a range of assays, we cannot say that for every combination. For example, we have not measured T cell function after cryopreservation in cervical or vaginal tissues. A related limitation is that the exploratory experiments shown in **Fig 1A-C** and **Fig 1E** were conducted with very small sample sizes. These experiments were meant to guide subsequent protocol development and narrow the range of possible procedures to the most promising. For this reason, we used these experiments to inform subsequent experiments, but did not perform statistical testing or draw firm conclusions from them.

Though our results suggest that cryopreservation of intact colorectal and cervicovaginal tissues should work well, there may be mucosal samples or assays other than those tested here where cryopreservation of intact tissue is not feasible or optimal. For example, we did not assess the effect of cryopreservation or vitrification on mucus production, quality, or function, so it is unknown whether mucus is adequately preserved by the protocols investigated here. In such cases, cryopreservation of cell suspensions, vitrification of tissues, or another protocol entirely may be better. To facilitate testing of vitrification for uses where cryopreservation is inadequate, a detailed vitrification protocol is included in **S1 Text** and at https://dx.doi.org/10.17504/protocols.io.p6adrae. A potential advantage of vitrification is that it avoids the need for a −80°C freezer and only requires a supply of liquid nitrogen, rather than both, as is the case for cryopreservation.

An important caveat with regard to vitrification is that it is possible that our protocol did not achieve true vitrification. Vitrification requires incredibly fast cooling rates and high concentrations of cryoprotective agents (in this case, DMSO and EG). The procedure used here may not have achieved adequate intracellular concentrations—due to slow permeation of the cryoprotective agents into the tissues—or sufficiently fast cooling. Further, the high concentration of the cryoprotective agents may itself have damaged cells in the tissue. A different procedure for vitrification might yield better results and more work could be done in this area.

Previous work on the preservation of the cervical tissues has been somewhat contradictory. In 2006, snap-frozen cervical tissue from surgical procedures was reported to have similar histology and susceptibility to HIV infection as fresh cervical tissue [5]. A second group has published similar data in terms of tissue viability (by MTT [3-(4, 5-dimethylthiazolyl-2)-2,5-diphenyltetrazolium bromide] assay) and histology for cryopreserved surgical tissue [3]. In contrast, cervical biopsies cryopreserved in 10% DMSO have been reported to support very little HIV replication [4]. Our data presented here suggest that fresh and cryopreserved ectocervical tissue explants are infected by HIV at similar rates. The discrepant results across these studies may owe to differences in (1) exact freezing and thawing procedures, (2) HIV strains, (3) age and other clinical conditions of the tissue donors, and (4) the uniformity of the biopsies (clinical research biopsies being more variable in size and more challenging to infect with HIV than the uniform explants taken from discarded surgical tissue used in this study). Further, the differences in infection rates of cryopreserved tissues may owe to the stringency of the HIV challenge. Our protocol, previously used to measure antibody-mediated protection against HIV infection [19], leads to infection of upwards of 90% of fresh explants. While this high level of infection is ideal for testing the protective efficacy of antibodies, it may overpower small changes in tissue viability.

There has been one previous reported attempt at preserving intact colorectal tissue. They found that snap freezing colorectal biopsies in liquid nitrogen, with or without a cryoprotectant, caused extensive tissue damage and reduced levels of HIV infection [2]; the tissue damage and reduction in infection were less severe after slow freezing with 7% DMSO. They also reported that 100% of fresh biopsies were infected after challenge with 1×10^4^ TCID_50_ of HIV_-1-BaL_, compared to 67% of cryopreserved, and 93% of those challenged with 100 TCID_50_, compared to 44% of cryopreserved. With our challenge dose of 1×10^4^ TCID_50_ of HIV_-1-BaL_ 90% of fresh explants and 74% of cryopreserved explants were infected, closely matching what was found in the previous study [2]. The reduction in efficiency of HIV infection in the colorectal tissue seen in both studies, as compared to ectocervical tissue, may indicate that colorectal tissues are less tolerant of cryopreservation than ectocervical or vaginal tissues.

The data presented in Figure 7 suggest that microbicide drug concentrations are lower in cryopreserved tissues than in fresh tissues, in particular for tenofovir diphosphate. However, small sample sizes and wide confidence intervals limit the conclusiveness of drug concentration results. One notable outlier in the drug concentration data was colorectal explants exposed to 0.759 nM dapivirine, where more than six times as much drug was measured in the cryopreserved explants as in the fresh ones. It is important to note that the absolute drug concentrations measured in this condition, where tissues were exposed to the lowest level of dapivirine used in the study, were very low, so this result may simply reflect noise in the measurements. Overall, our data do not rule out the feasibility of measuring drug concentrations in cryopreserved tissues, but indicate the need to test the effect of cryopreservation individually for each drug of interest, as the effect may vary strongly by chemical compound.

In this article, we used non-inferiority testing, because it is more informative than determining equivalence by p > 0.05 in a standard statistical test. Our studies were not designed with this analysis plan in mind and, due to practical constraints, our studies have small sample sizes with limited power to detect non-inferiority. While we did find the metabolic activity of cryopreserved and vitrified vaginal tissues to meet our definition of non-inferior, this was not the case for HIV infection assays. Although the infection hazard ratios were very similar between fresh and preserved tissues, the confidence intervals were too wide to meet our 20% threshold of non-inferiority. This may be a result of the small sample sizes or of a difference between fresh and preserved that we did not have the power to discern.

The experiments described in this article were conducted using samples stored in liquid nitrogen vapor for between a few minutes and a few months, so we have not formally demonstrated that sample quality is maintained with long-term storage. However, as also discussed in our previous paper [1], storage at liquid nitrogen temperatures should theoretically maintain samples unchanged indefinitely, and this has been shown experimentally for timescales of years [44,45]. Therefore, we believe that samples stored according to our protocols will remain stable indefinitely.

This is the last of a series of papers reporting our studies designed to provide guidance on cryopreserving viable cells and tissues from mucosal sites of sexual pathogen transmission [1,17,18]. In the step-by-step protocols in **S1 Text** and the Supplement to [1], we aim to provide clear procedures to ensure that cryopreservation can be carried out in the best known manner. Our data support the use of immunological assays and *ex vivo* HIV infections with cryopreserved mucosal cell and tissue specimens.

## Acknowledgments

We wish to dedicate this article to the memory of coauthor Charlene Dezzutti, much missed friend and scientist, taken from us too soon. None of this research would have been possible without the generous participation of the study volunteers. Thank you to Katharine Westerberg for assistance with processing vaginal tissues. We are grateful to Dr. Christina Ochsenbauer for sharing her HIV constructs and Greg Mize for making the nanoluciferase-secreting virus. We also thank Dr. Peter Hunt, Dr. Ma Somsouk, Ms. Becky Hoh and the San Francisco General Hospital SCOPE team for recruiting participants and obtaining gastrointestinal biopsies. Thank you to the Fred Hutch Experimental Histopathology Core for performing H&E stains. We are grateful to Dr. Patricia D’Souza (DAIDS, NIH) for support and encouragement. We thank Dr. Sarah Holte (Statistical Core, Center for AIDS Research, University of Washington; funded by NIH grant AI027757) for statistics consultation and Veronica A. Davé for critically reviewing the manuscript. We acknowledge Dr. Robert C. Hackman for helpfully reviewing the H&E-stained vaginal tissue sections. The University of Pittsburgh Medical Center Hillman Cancer Center and Tissue and Research Pathology/Health Sciences Tissue Bank shared resource, which is supported in part by award P30CA047904 provided surgical remainders for the work in Dr. Charlene Dezzutti’s laboratory. The NIH AIDS Reagent Program, Division of AIDS, NIAID, NIH provided tenofovir; International Partnership for Microbicides provided dapivirine; and Merck & Co., Inc., Kenilworth, NJ USA provided MK-2018.

## Supporting information

**S1 Fig.**
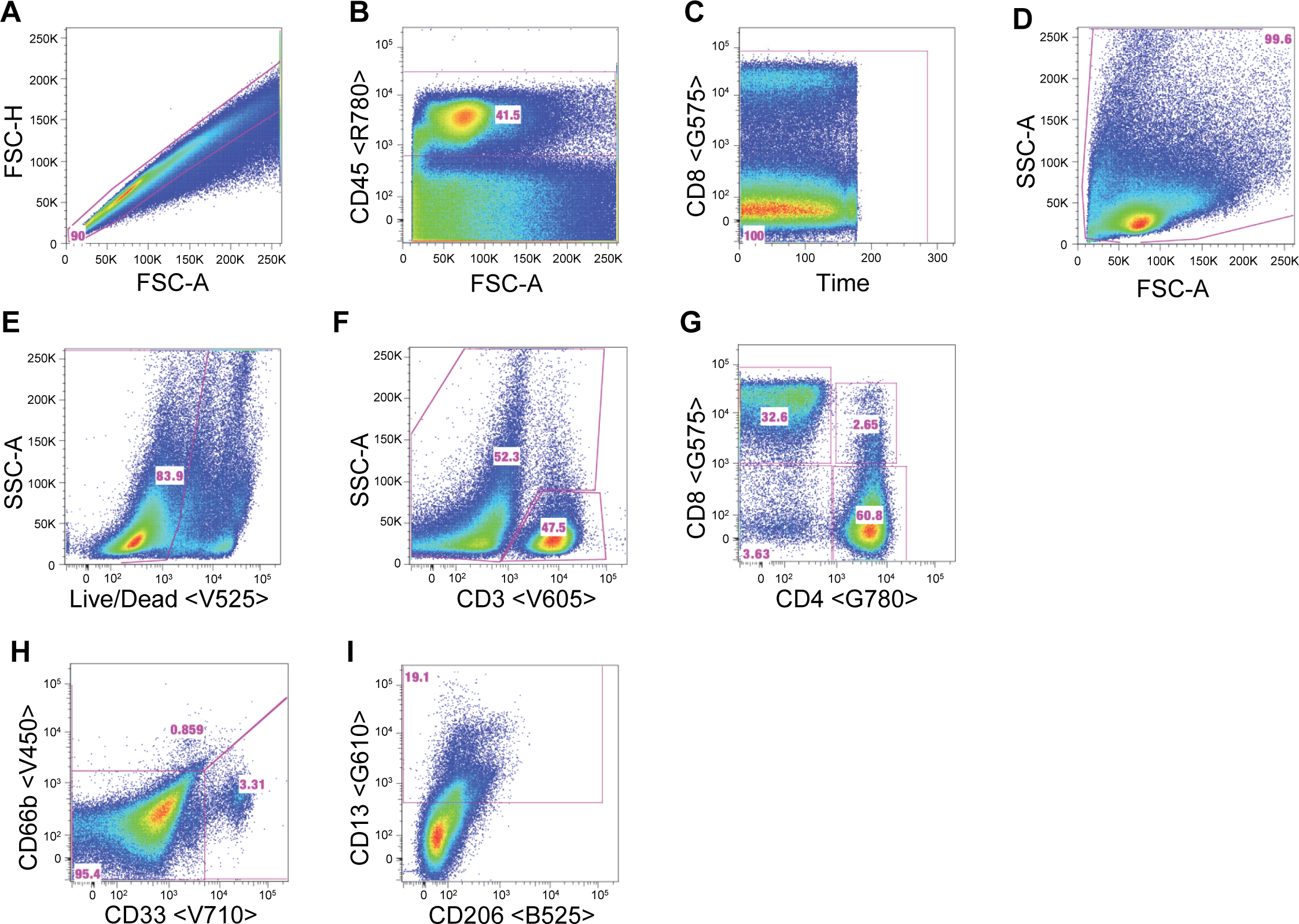
Flow cytometric gating of cryopreserved and vitrified colorectal cells. **A**, Exclusion of doublets. **B**, Selection of CD45-expressing leukocytes. **C**, Time gate for removal of spurious events caused by flow disruptions. **D**, Exclusion of debris by forward and side scatter. **E**, Selection of live cells by exclusion of stained dead cells. **F**, Separation of cells into CD3-expressing T cells and non-T cells. **G**, Separation of T cells into CD4 and CD8 subsets. **H**, Identification of CD66b-expressing granulocytes and CD33-expressing myeloid cells from the non-T cells. **I**, Identification of CD13-expressing monocytes from the non-T cells. “FSC” and “SSC” refer to forward and side scatter, with “-A” indicating area and “-H” indicating height. Fluorescent detectors are indicated inside angle brackets, with the first letter indicating the laser (red, green, violet, or blue) and the three digits indicating the middle of the bandpass filter for the detector (in nanometers).

**S2 Fig.**
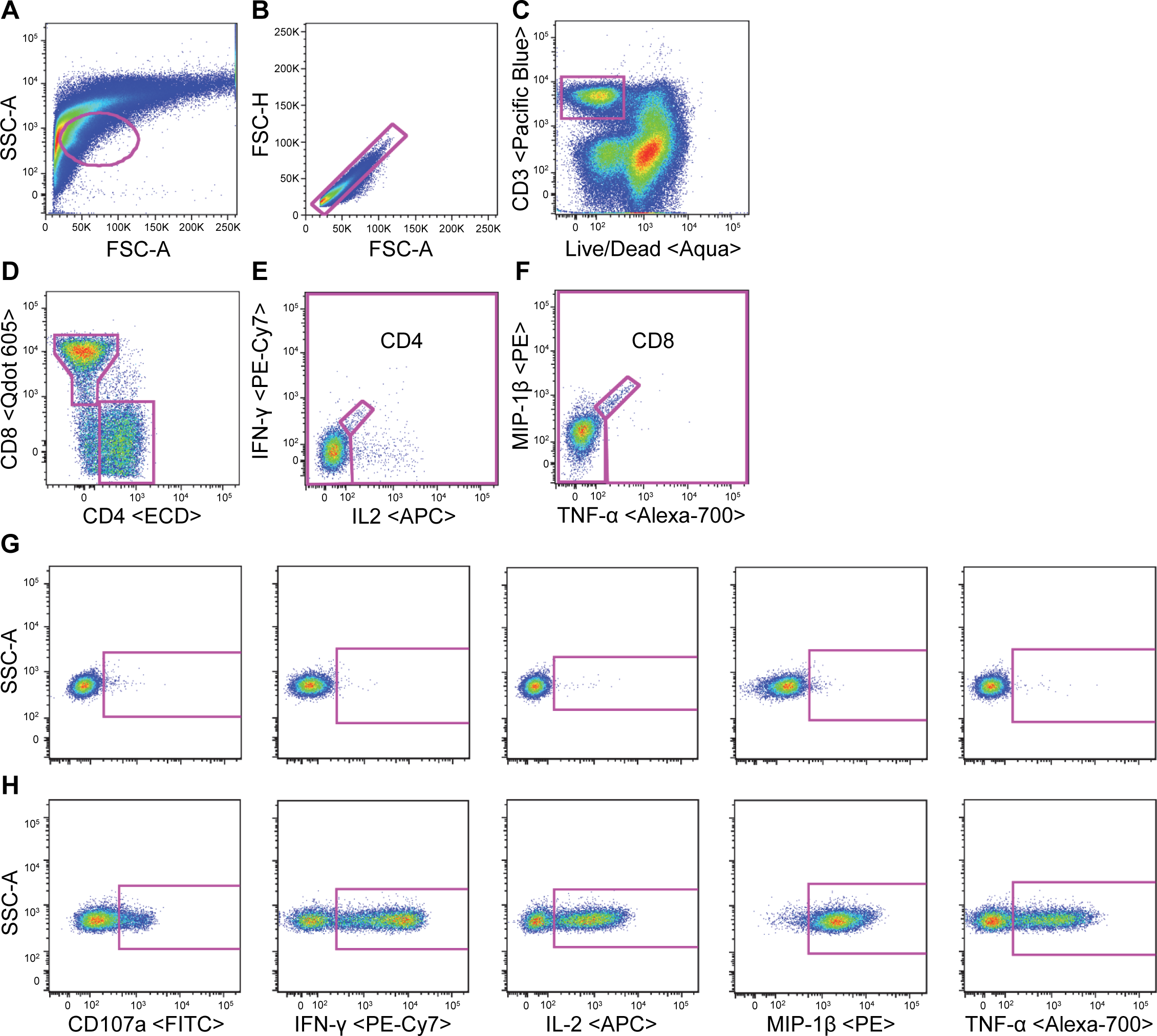
Flow cytometric gating of ex vivo stimulated and intracellular stained colorectal cells. **A**, Selection of lymphocyte-like cells by forward and side scatter. **B**, Exclusion of doublets. **C**, Identification of live CD3+ T cells. **D**, Separation of T cells into CD4 and CD8 subsets. **E**, Exclusion of non-specific staining from CD4 cells. All events are included in this gate except for those inside the diagonal box. **F**, Exclusion of non-specific staining from CD8 cells. **G**, Identification of cytokine- or CD107a-expressing cells in the unstimulated condition. **H**, Identification of cytokine- or CD107a-expressing cells in the PMA/ionomycin-stimulated condition. “FSC” and “SSC” refer to forward and side scatter, with “-A” indicating area and “-H” indicating height. Cytokines measured were interferon-γ (IFN- γ), interleukin-2 (IL-2), macrophage inflammatory protein (MIP)-1β, and tumor necrosis factor-α (TNF-α). “APC” indicates allophycocyanin.

**S1 Table. Antibody source table.** Reagent information for all antibodies.

**S1 Text. Detailed, step-by-step protocols for cryopreservation and vitrification of mucosal tissues.**

**S1 File. Complete statistical tables.**

**S2 File. Analysis code and data.**

